# Cell cycle lengths of stem cells and their lineage from cellular demography

**DOI:** 10.1101/2020.11.18.388314

**Authors:** Purna Gadre, Nitin Nitsure, Debasmita Mazumdar, Samir Gupta, Krishanu Ray

**Affiliations:** Department of Biological Sciences, Tata Institute of Fundamental Research, Mumbai, Maharashtra 400005, India; School of Mathematics, Tata Institute of Fundamental Research, Mumbai, Maharashtra 400005, India

**Keywords:** Germline-stem-cells, Transit-Amplification, Cdc25/String, Cell cycle, demographic model

## Abstract

Adult stem cells and their transit-amplifying (TA) progeny dynamically alter their proliferation rates to maintain tissue homeostasis. To test how the division rates of stem cell and TA cells affect tissue growth and differentiation, we developed a computation strategy which estimates the average cell cycle lengths/lifespans of germline stem cells (GSCs) and their TA progeny from cellular demography. Analysis of the wild-type data from *Drosophila* testis using this method indicated anomalous changes in lifespans during the germline transit-amplification with a nearly 1.3-fold increase after the first division and about a 2-fold decrease in the subsequent stage. Genetic perturbations altering the cell cycle rates of GSC and its immediate daughter, the gonialblast (GB), proportionately changed the rates of subsequent TA divisions. Notably, a nearly 2-fold increase or decrease in the total TA duration did not alter the induction of meiosis after four mitotic cycles. Altogether, these results suggest that the rates of GSC and GB divisions can adjust the rates of subsequent divisions and the onset of differentiation.

**Significance Statement:** Dynamic regulation of the proliferation rate of stem cells and their transit-amplifying daughters maintains tissue homeostasis in different conditions such as tissue regeneration, aging, and hormonal imbalance. Previous studies suggested that a molecular clock in the stem cell progeny determines the timing of differentiation. This work shows that alterations of the rates of stem cell divisions, as well as that of its progeny, could override the differentiation clock in the *Drosophila* testis, and highlights a possible mechanism of fine-tuning the transit-amplification program under different conditions such as tissue damage, aging, and hormonal inputs. Also, the method developed for this study could be adapted to estimate lineage expansion plasticity from demographic changes in other systems.

**Highlights:**

- Determination of cellular lifespan during transit-amplification from demography
- Lifespans of Drosophila male germline cells changes anomalously during the TA
- Lifespan changes of germline stem cells readjust that of the progeny cells
- Anomalous lifespan expansion midway through TA precedes the Bam onset

## Introduction

Many adult stem cells produce progenitors, which undergo transit-amplifying (TA) divisions, before terminal differentiation. Hormonal stimulation (1, 2), tissue damage (3, 4), aging (5–7), etc., alter the division rates of stem cells and their progeny. Previous studies have shown that the TA cells pass through a continuum of transcriptomic states, setting the timing of differentiation, independent of the TA cell cycle rates (8–10). Coordination of this autonomous differentiation clock with the rates of TA divisions is essential for tissue homeostasis, defects in which can lead to cancer or other disorders (11, 12). Despite its importance, it is unclear whether the rates of stem cell and TA divisions influence the differentiation clock.

*Drosophila* spermatogenesis provides an ideal model system to study the regulation of TA divisions. Accumulation of a translational repressor, Bag-of-marbles (Bam), to an optimum level arrests the TA divisions after the 4^th^ round, suggesting that the bam expression and degradation kinetics could set the differentiation clock (10). It was also evident that, slowing down the 3^rd^ and 4^th^ TA divisions (of the 4- and 8-cell stages, respectively) could induce premature germline differentiation after the 3^rd^ round, but the effect was not fully penetrant (10). Several cysts concluded the TA and meiosis in a wild-type-like manner despite the modification. This observation also suggests that the differentiation clock can adjust to accommodate changes in the rates of TA divisions. Furthermore, perturbing the rates of GSC divisions and early TA divisions did not affect the differentiation at the 16-cell stage (13). Together, all these investigations highlighted that, up to a limited extent, the rates of TA divisions could influence the differentiation clock. However, we still lack clarity regarding the quantitative limits of this readjustment.

To resolve this issue, one requires an estimate of how the cell cycle lengths of GSCs and TA cells change under different conditions. Previous studies inferred the changes in the proliferation rates of the GSCs and TA cells by enumerating phospho-histone3/BrdU stained clusters (13–16) and performing BrdU/EdU pulse-chase analysis (10, 17). Although these methods presented a comparative measure to examine how factors regulate the GSC and TA pool, they failed to quantitate the cell cycle lengths of the GSC and TA stages. Recently, the time between two successive GSC divisions (inter-division lifespan) was estimated using time-lapse imaging of isolated testes for up to 19 hours (18, 19). While time-lapse imaging measures the exact length of the cell cycle, it is tedious for a multi-factor manipulation of the cell cycle rates. Moreover, long-term exposure of cells to light while imaging increases phototoxicity due to ROS generation, which might alter the very rate it is supposed to measure.

Therefore, we devised an optimized computation strategy to estimate the cell cycle lengths, hereafter referred to as the ‘lifespans’, of the GSCs and TA stages using five parameters: 1) Size of GSCs and TA population, 2) GSC mitotic indices, 3) GSC M-phase duration, 4) Germ cell death frequency and 5) Persistence time of a dead cyst. This method restricted the requirement for time-lapse imaging and enabled us to examine the effects of a range of genetic perturbations on the GSC and TA division rates. Using this method, we probed the correlation between the GSC division rates and the onset of the differentiation program. The results suggest that altering the rates of stem cell divisions and early TA divisions affects those of the subsequent divisions and readjusts the differentiation clock to such an extent that the scale and relative pattern of the germline amplification remains unaltered.

### Formulation of the equation for empirical estimation of the cell lifespans of the TA stages

In wild-type testes, the GSCs surround the stem cell niche, termed as the hub (Fig. 1A and B). Each TA division displaces the resultant cyst further away from the hub. The GSCs and the TA stages can be visualized by immunostaining the testes for Vasa (which marks the germline cells), Armadillo (labels the hub and the cyst perimeter), and MAb1b1/Hts1 (labels spectrosomes and fusomes; Fig. 1B). To compute the lifespans of the GSCs and TA cells, we assumed the following:

1. The TA population in the adult testis is in a steady-state (Fig. 1E, discussed in supplement section 6).
2. The lifespan of each stage (GSC and TA stages) remains invariable (see supplement section 6).
3. The cell cycle phases at each stage (GSC and TA stages) are uniformly distributed across the cells in that stage (see supplement section 6).

**Fig. 1.**
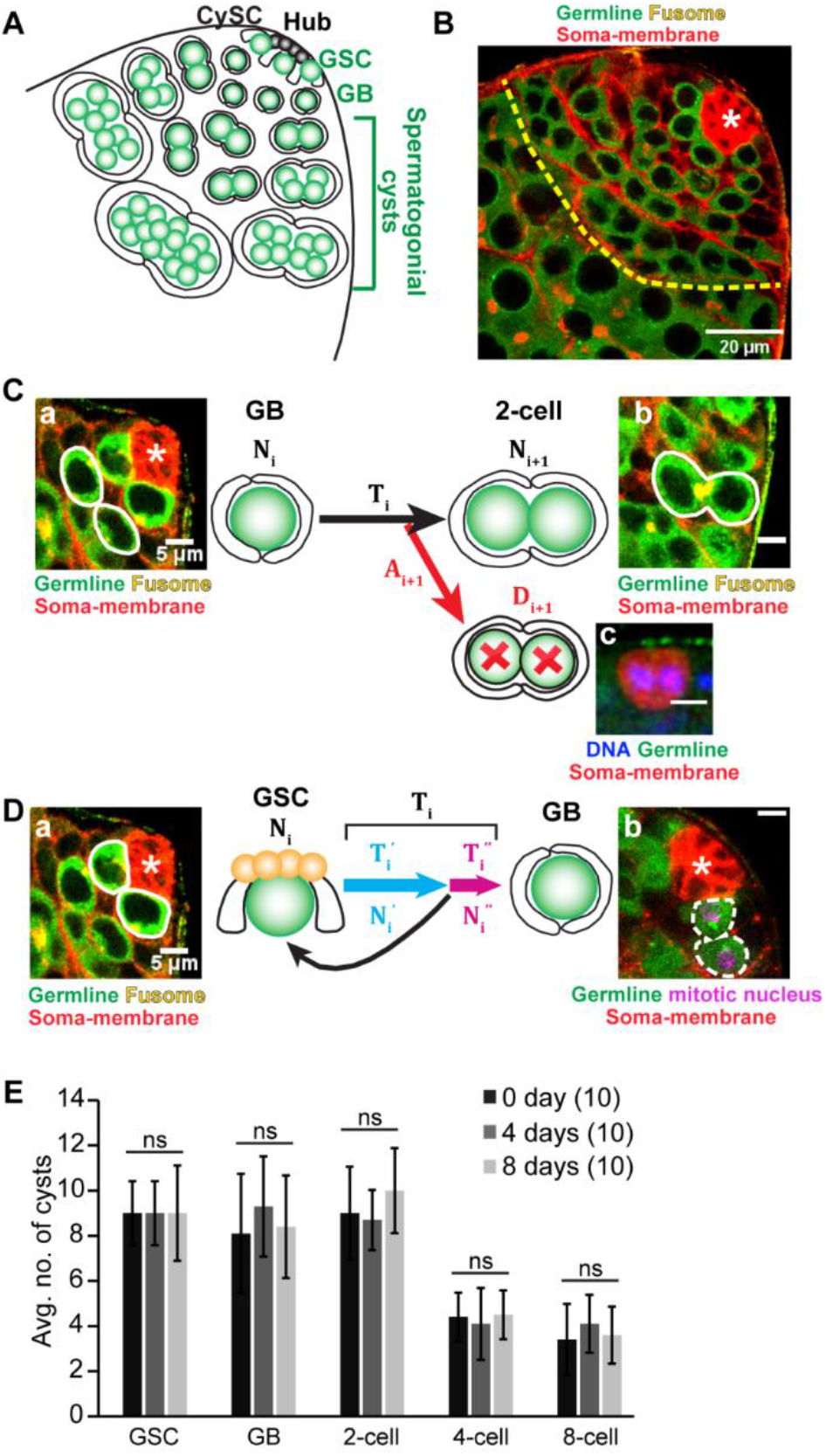
A) Schematic illustrates the process of transit-amplification during early spermatogenesis. Glossary: GSC – Germline Stem Cell, CySC – Cyst Stem cell, GB – Gonialblast. B) The apical tip of wild-type (*CantonS*) testis stained for Vasa (Green), Armadillo (Red), and Adducin/Hts (Red). (Scale bars ~20μm). C) Schematic describes the sequential method (equation (1)) of time estimation. *T_i_* denotes the time taken for a GB to 2-cell cyst transition (GB lifespan). *N_i_* and *N*_*i*+1_ denote the number of GBs (a) and 2-cell cysts (b), respectively. *D*_*i*+1_ denotes the number of 2-cell dead cysts, depicted by the Lysotracker-positive 2-cell cyst (c). *A*_*i*+1_ denotes the persistence time of a dead cyst. (Scale bars ~5μm). D) Schematic describes the individual method (equation (2)) of time estimation. 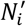 and 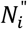 denote the number of GSCs in G1/S/G2 phases (a) and M-phase (b), respectively. 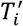 and 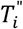 denote the duration of G1+S+G2 phases and that of the M-phase, respectively. *T_i_* denotes the GSC lifespan. (Scale bars ~ 5μm). E) Histograms show the relative stage-specific distribution profile of cysts (average ± S.D.) in *CantonS* adults aged for 0-, 4- and 8-days after emergence from the pupal case (eclosion) at 29°C. (n=10 for each group. Kruskal–Wallis test, p>0.05 for all stages).

In such a closed system, at steady-state, the relative population of a stage 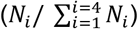 is equal to its relative lifespan 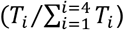, if no germline cysts are lost due to death. Here, *i* represents the TA stage (taking four values corresponding to the four TA stages: GB, 2-cell, 4-cell, 8-cell), *N_i_* represents the average number of cysts at stage *i*, and *T_i_* denotes the lifespan of the stage *i*, respectively (Supplemental Methods section 1-7 for detailed reasoning). For two successive stages *i* and *i* + 1, this relationship can be expressed as,

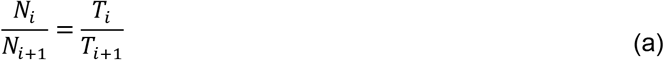

### Adaptation of the computation strategy to account for the germ cell death

Reportedly, a small number of cysts undergo germ-cell death (GCD) at every TA stage (20–22). Hence, only a fraction of cysts successfully transitions to the next stage. In our model, we assumed that, for a transition of a cyst from stage *i* to *i* + 1, the GCD occurs after stage i cysts complete the cell cycle (*i. e*., concluding the G1, S, G2, and M), and before they enter the next cell cycle of stage *i* + 1. To account for this loss, we defined the survival probability (*s*_*i*+1_) as the possibility of a successful transition from stages *i* to *i* + 1 (see supplemental Methods sections 7, 8, and 9). Incorporating this survival probability (*s*_*i*+1_) in equation (a) gives,

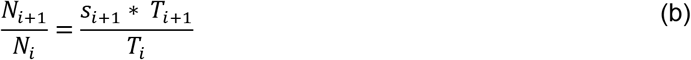

*s*_*i*+1_ can be calculated by using the same logic as equation (b): the fraction of dead spermatogonial cysts (*D*_*i*+1_/*N_i_*) during the transition from stages *i* to *i* + 1 is equal to the product of the probability of GCD (*d*_*i*+1_ = 1 – *s*_*i*+1_ and the relative persistence time of the dying cysts (*A*_*i*+1_/*T_i_*). *D*_*i*+1_ denotes the average number of cysts that died during the transition from stage *i* to stage *i* + 1, and *A*_*i*+1_ denotes the time for which a dead cyst remains visible in the testis (persistence time). This relationship is expressed as:

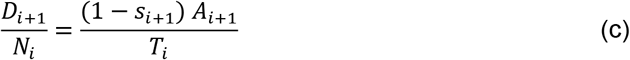

After a rearrangement of equation (c) we get,

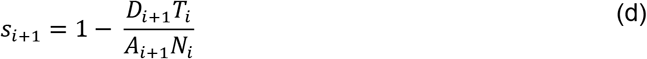

### Solving the equations to obtain the lifespans

#### 1. Sequential estimation method (Fig. 1C)

(b) and (d) give two equations in the three unknowns *T_i_, T*_*i*+1_ and *s*_*i*+1_.

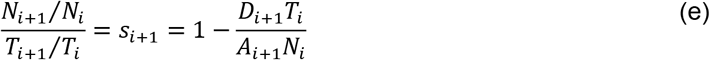

Eliminating *s*_*i*+1_ and solving for *T*_*i*+1_ gives

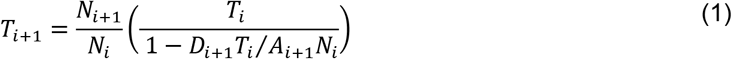

#### 2. Individual estimation method (Fig. 1D)

Alternatively, the cyst population of stage *i* can be divided into two sub-stages: the cyst population in G1/ S/ G2-phases and M-phase. We denote these two stages as stage *i′*, and *i″*, respectively. As mentioned earlier, our assumption implies that no deaths occur during the phases G1, S, G2, and M of the germline cell cycle. As the sub-stage *i′* is by definition made up of the phases G1, S, and G2, and as the sub-stage *i″* is the corresponding M-phase, the assumption implies that any spermatogonial cyst in sub-stage has a 100% probability of transitioning to the sub-stage *i″*. Consequently, we have,

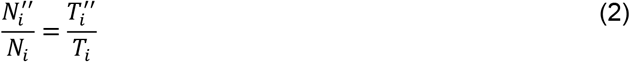

where 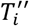 and *T_i_* are M-phase length and the lifespan of the stage *i*, respectively. 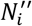 and *N_i_* are the number of cysts of stage *i* in M-phase and the total number of cysts of stage *i*, respectively. The above relationship shows that the fraction of cysts of stage *i* in M-phase (Mitotic index, 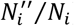) is equal to the length of the M-phase relative to the total lifespan of stage *i* (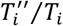; see supplement Methods section 10).

These methods provide a population average estimate of the inter-division lifespan for a GSC or the cell cycle length/ lifespan of a TA stage. The accuracy of the estimate depends on the sample size of each parameter and penetrance of the genetic/ environmental perturbation. Nevertheless, the method is internally consistent which would allow comparative quantitative analysis to examine the effect of a perturbation.

## Results

### The TA population in the adult testis is in a steady-state

To explore the germline population dynamics in the adult testis, we enumerated the GSCs and TA stage-wise distribution of the cysts from 1) freshly emerged (0-day), 2) 4-day old, and 3) 8-day old wild-type males. The cyst distribution remained invariant, demonstrating that the germline TA population is in a steady state until 8-days post eclosion (Fig. 1E).

### GSCs and TA stage cysts take nearly equal time to complete their M-phases

Phospho-Histone-3 (pH3) staining in fixed preparation efficiently identified the mitotic stages from prophase to telophase (Fig 2A, B; (23)). To determine the M-phase length of the GSCs and cysts in live preparations, we collected time-lapse images from *nosGal4vp16>UAS-Histone-RFP; Jupiter-GFP* (*nos>His-RFP; Jup^PT^*) testes *ex vivo. Histone-RFP* expression labeled the nuclei of GSC and its progeny cells (Fig. 2A, B, D). Jupiter-GFP marked the microtubules (24), and the morphology of microtubules allowed us to identify the onset and termination of M-phase as described previously (25). The appearance of two centrosome-associated microtubule clusters was considered the beginning of prophase (arrowhead, Fig. 2D-a; (23, 26)). Metaphase was identified by the characteristic spindle formation and chromatin alignment at the cell equator (arrow, Fig. 2D-c). The spindle then resolved through anaphase (Fig. 2D-d) and telophase (Fig. 2D-e). A visible increase in the nuclear size during the time-lapse series was considered as the end of telophase (Fig. 2D-f). The GSC M-phase period varied from 50 to 80 minutes (N = 8, Fig. 2E). We used the median of this dataset (67.5) for our calculations. The M-phase durations remained invariant in the GB, 2-cell, 4-cell, and 8-cell cysts (Fig. S2, Fig. 2E).

**Fig. 2.**
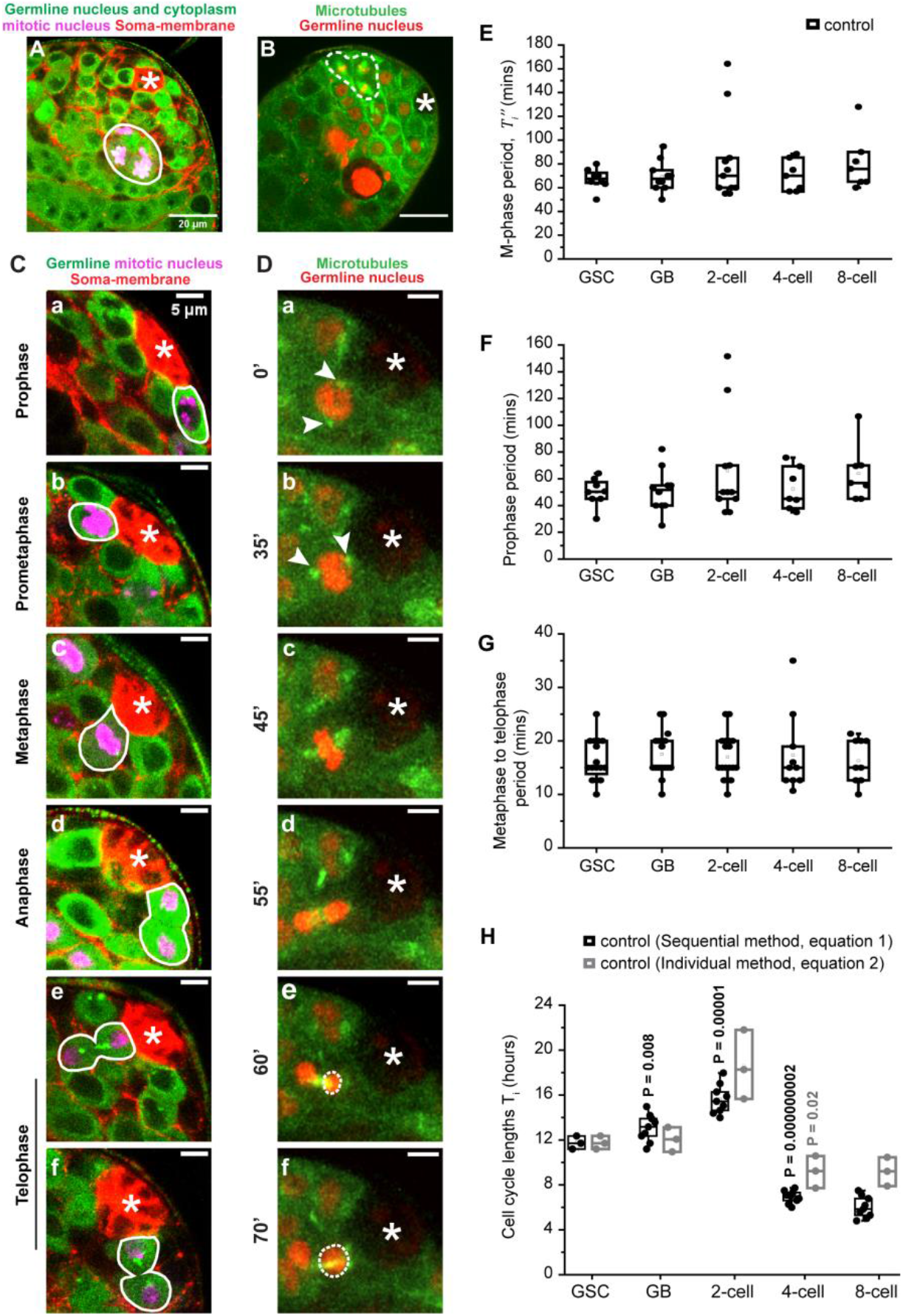
A) Apical tip of *nos>His-RFP* (Green) testis stained for Vasa (Green), Armadillo (Red) and pH3 (Magenta). The white boundary marks a pH3-positive 4-cell cyst. B) A snapshot from the time-lapse imaging of the apical tip of *Jupiter^PT^*(G reen); *nos>His-RFP*(Red) testis. Dotted white boundaries mark a 4-cell cyst in metaphase (Scale bars ~20μm) C) Adult testis tip stained for Vasa (Green), Armadillo (Red), and pH3 (Magenta) showing a GSC in prophase (a), pro-metaphase (b), metaphase (c), anaphase (d) and telophase (e, f) marked by white boundaries. D) Montage of a time-lapse image of a GSC (*Jupiter^PT^*(Green); *nos>His-RFP*(Red)) undergoing prophase (a), prometaphase (b), metaphase (c), anaphase (d) and telophase (e, f). Arrowheads mark the position of the separated centrosomes. White dashed circles show the increase in the nuclear size. Time intervals in minutes are indicated next to the image panels. (Scale bars ~ 5μm). In all images, asterisk marks the hub. C-E) Box plots show the duration of M-phase (E), prophase (F), and metaphase to telophase (G) in GSCs and TA stages in control (*nos>His-RFP; Jupiter^PT^*). (Mann Whitney-U test). H) Box plot shows the lifespans of GSCs and TA stages in control using equation (1) and equation (2). (Students T-test).

### The GSCs divide every 12 hours

In wild-type testis, no GCD was recorded in GSCs (N=54) (27). Therefore, we calculated the average time between two successive GSC divisions (inter-division lifespan) directly from the M-phase period (Fig. 2E) and GSC mitotic index (Table S1) using equation (2). The average GSC lifespan in control (*nos>EGFP*) background was estimated to be 11.7 ± 0.6 hours (Fig. 2H), which falls within the reported range (19).

### Cellular lifespans increase for the first two TA divisions and then contract by nearly 2-folds

To estimate the lifespans of the spermatogonial TA stages, we used equation (1). We used the consolidated, stage-wise cyst counts (*N_i_* and *N*_*i*+1_’s; Table S2) obtained from the previously published literature (13), and enumerated the Lysotracker-positive cysts in phase-I from the same genetic background to obtain *D*_*i*+1_’s (Table S3). The stage-wise phase-I persistence time (*A*_*i*+1_’s) obtained from the time-lapse images (Fig. S1A, Table S4) varied substantially across samples, ranging from 1 to 4 hours (Table S4). A limited simulation suggested that for values between 1 and 5 hours, the lifespan estimates (*T_i_*’s) remain fairly unaltered (Fig. S1B). This justifies our assumption of the average of the observed values of phase-I persistence time as a constant value for *A* (Table S4).

The estimates suggested that the lifespan of GBs (13.6 ± 1.2 hours) is about 16% longer as compared to that of the GSCs, and the 2-cell lifespan (17.6 ± 1.6 hours) is further prolonged by approximately 29% (Fig. 2H). Subsequently, the lifespans of 4-cell cysts contracted by about 54% (8.1 ± 0.6 hours), and remained at a similar level during the 8-cell stage (7.3 ± 1.1 hours; Fig. 2H). These observations were consistent with a previous report suggesting that the 4- and 8-cell cysts take more than 7 hours to complete their cell cycle (10).

To confirm these estimates, we sought to calculate the lifespans by an alternate method using equation (2), which utilizes the stage-wise mitotic indices (Table S1) and the M-phase durations (Fig. 2E). Consistent with the results obtained using equation (1), these estimates indicated similar changes in the lifespans at the 2- and 4-cell stages (Fig. 2H).

Together, these analyses suggest that the TA lifespans increase after GB and then shrink by about 2-folds for the next two divisions. Hence, contrary to the earlier assumptions, we find that the transit-amplification is not a uniform process. These methods also indicated that the TA lasts of nearly 47 hours (Table S5), which is close to an earlier estimate (28). A recent study in the *Drosophila* ovary suggested anomalous alterations in the cell cycle structure during the TA stages, although the authors could not quantify the lifespans, (29). Also, developmentally regulated anomalous TA divisions were observed in the *Drosophila* type II neuroblasts lineage, in which the first daughter divides after 6.6 hours and the matured late-stage daughters divide every 2-3 hours (30). Together, these results suggest that the non-uniformity of the TA rates could be a recurrent theme in the stem cell systems.

### Autonomous disruptions of G1/S or G2/M transitions in GSCs and GBs prolongs the lifespans of all the TA stages

Next, to understand the impact of changes in the lifespan of stem cells, we perturbed the G1 and G2 phases in GSCs and early TA cells. In *Drosophila* epithelial cells, Cyclin E promotes G1/S transition (31), and *String/CDC25* (*stg*), via Cdk1 activation, induces the G2/M transition (32). We showed earlier that RNAi knockdown of *cycE, stg*, and *cdk1* in the GSC and GB does not alter the transit-amplification program in testis (13). Therefore, we probed the effects of these perturbations on the lifespans of the GSC and TA stages.

We reasoned that if the TA divisions are autonomously regulated independent of the GSC, the RNAi mediated knockdown of *cycE* or *cdk1* will increase the lifespans of only GSCs and GBs without affecting that of the subsequent TA stages. Time-lapse imaging suggested that the GSC M-phase duration is significantly altered only in the *nos>cdk1^dsRNA^* background (Fig. 3B-a). Whereas, computation of the lifespans from enumeration of the *D*_*i*+1_’s (Table S3), the GSC mitotic indices (Table S1), and the *N_i_*’s (Table S2; (13)) in the *nos>cycE^dsRNA^* and *nos>cdk1^dsRNA^* backgrounds, revealed more than a 2-fold increase in the GSC and GB lifespans, due to the *cycE* RNAi (*T_GSC_* = 24.7 ± 3.2 hours, *T_GB_* = 21.5 ± 2.7 hours; Fig. 3E) and the *cdk1* RNAi (*T_GSC_* = 33.3 ± 3 hours, *T_GB_* = 26.1 ± 2.9 hours; Fig. 3F), respectively (Table S5). A similar estimation was obtained using the eq-2 (Table S6). Contrary to the expectation, both the RNAi perturbations also prolonged the lifespans of 2-, 4-, and 8-cell stages by a similar margin (Fig. 3E, F), suggesting that the rates of the GSC/GB divisions could influence those of the subsequent TA divisions.

**Fig. 3.**
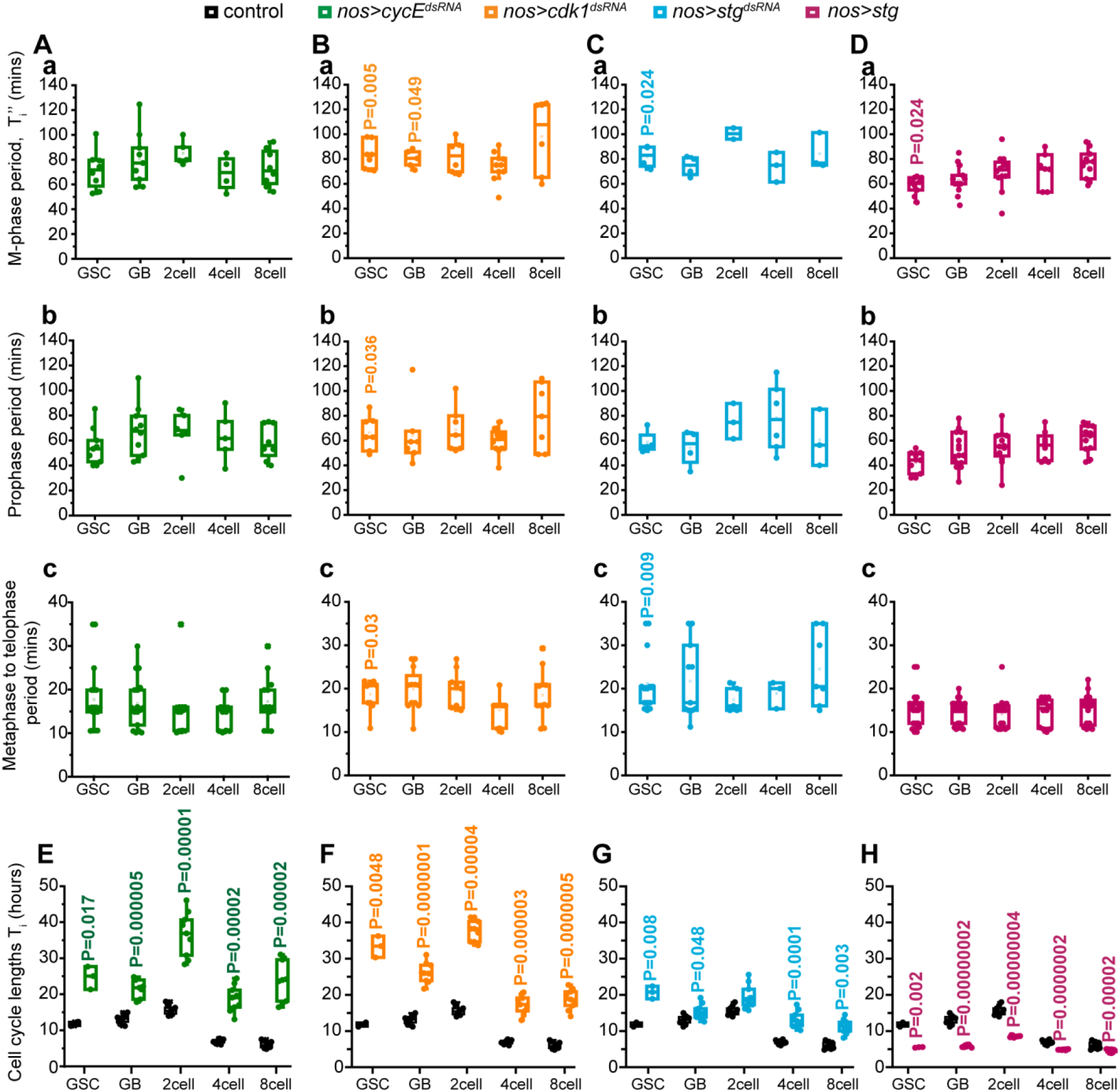
A-D) Box plots show the duration of M-phase (a), prophase (b), and metaphase to telophase (c) in GSCs and TA stages in *nos> His-RFP; Jupiter^PT^* background overexpressing *cycE^dsRNA^* (A), *cdk1^dsRNA^* (B), *stg^dsRNA^* (C), and *stg* (D). (P values calculated with respect to control in Fig. 2 using Mann Whitney-U test). E-H) Lifespans estimations in *nos>cycE^dsRNA^* (E), *nos>cdk1^dsRNA^* (F), *nos>stg^dsRNA^* (G), and *nos>stg* (H) backgrounds (Mann Whitney-U test for 2cells in (F), otherwise Students T-test).

To test this conjecture, we perturbed the cell cycle using Cdc25/string RNAi, which is predominantly expressed in the GSCs and GBs (33). As expected, the *nos>stg^dsRNA^* expression significantly increased the GSC M-phase period (Fig. 3C-a), as well as the lifespans of the GSCs (20.7 ± 1.8 hours) and GBs (15.4 ± 2.1 hours; Fig. 3G). Also, the lifespans of the 4-cell (13.3 ± 2.3 hours) and 8-cell (11.2 ± 2 hours) stages were significantly higher in *nos>stg^dsRNA^* background (Fig. 3G). Together, these results suggested that slowing down the cell cycle rates of GSCs and early TA stages have a feed-forward effect and prolong the subsequent divisions. Inexplicably, though the estimates of 2-cell lifespans, obtained using eq-1 and eq-2, respectively, in the stg RNAi background deferred by a significant margin. These results also suggested that the TA durations can be extended by more than 2 folds without affecting the homeostasis (Table S5).

### Ectopic Stg overexpression speeds up the entire transit-amplification program

Ectopic stg overexpression at the 4 and 8-cell stages was shown to shorten the cell cycle lengths (10). The *nos>stg* overexpression significantly shortened the GSC M-phase period (Fig. 3D-a), and significantly increased the GSC mitotic index (Table S1) (13). Expectedly, the stg overexpression shortened the lifespans of the GSC (5.8 ± 0.2 hours) and GB (5.8 ± 0.3 hours) by approximately 2-fold (Fig 3H). Furthermore, the lifespans of 2-cell (8.5 ± 0.2 hours), 4-cell (4.9 ± 0.1 hours), and 8-cell (4.6 ± 0.5 hours) stages were also shortened by approximately 2-folds (Fig. 3H).

Together, these results suggested that perturbing the cell cycle lengths of GSCs and early TA stages affects the lengths of all the TA divisions and consequently, the entire transit-amplification program. We noted that the anomalous pattern of the TA lifespans was conserved in all these instances (Fig 4A). Remarkably, altering of the GSC lifespans readjusted the lifespans of subsequent stages (Fig 4B). Hence, the stem cell lifespan appeared to set the duration of transit-amplification and the onset of differentiation.

**Fig. 4.**
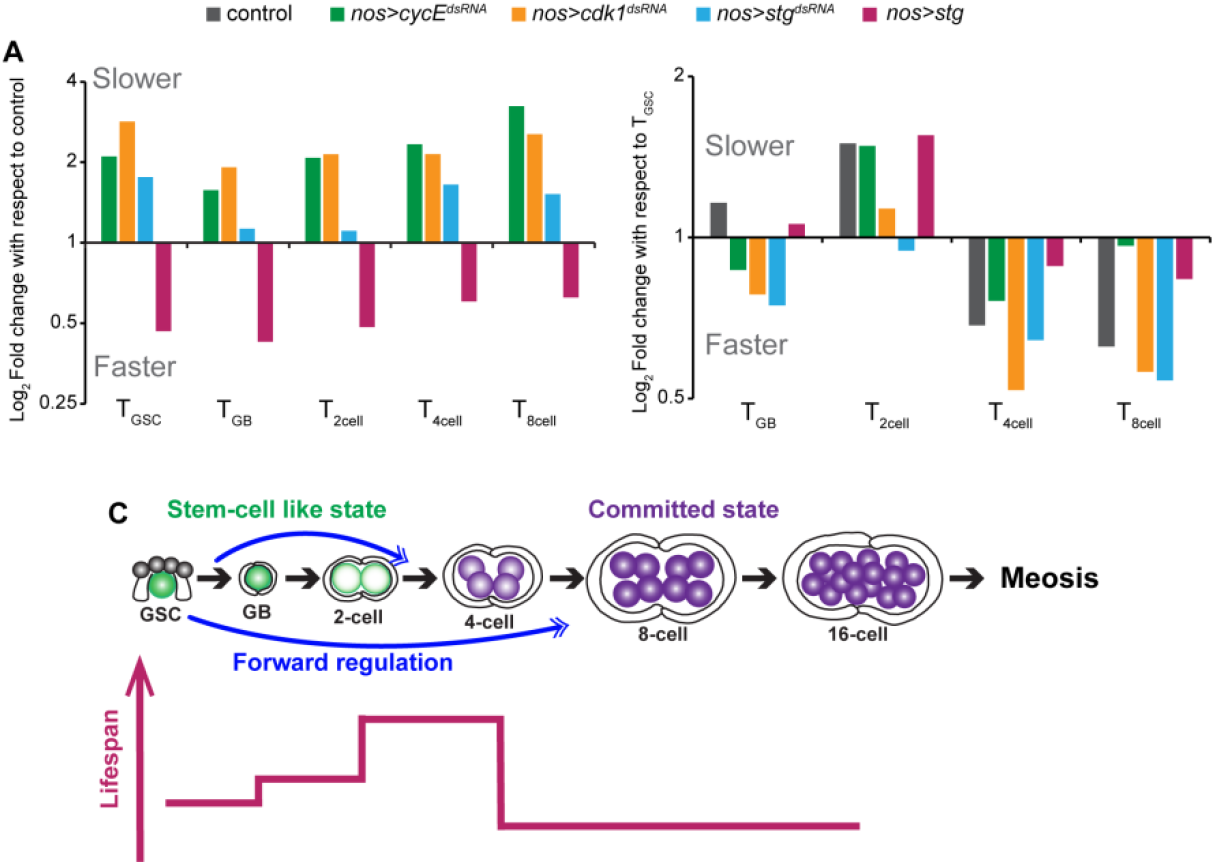
A) Log_2_ fold change in lifespans of GSCs and TA stages in different genetic backgrounds relative to control. B) Log_2_ fold change in lifespans of TA stages in different genetic backgrounds relative to the GSC lifespans in respective backgrounds. Note that in control, the lifespans increase to about 1.5 fold and decrease by about 1.5 folds in 4- and 8-cell stages. C) Schematic illustrates the germline TA lineage in the *Drosophila* testis. The lifespans undergo anomalous alteration with the TA progression. The 2-cell to 4-cell transition presumably signifies the shift from a stem-cell-like to a differentiated cell-like state. Stem cells send a feed-forward signal to regulate the rates of TA divisions, thus fine-tuning the timing of entry into meiosis.

## Discussion

In the *Drosophila* testis, the GSCs can be easily identified due to the floral arrangement around the hub (Fig. 1A, B); whereas, the spermatogonial cysts are more abundant and tightly packed with no specific spatial marker for identification in live tissue. Hence, the time-lapse measurement of the cell cycle length was limited to the GSCs (18, 19). We devised a computation strategy that estimates the average division rates of the GSC and TA at steady-state. This method requires time-lapse measurement of the M-phase period of GSCs and the persistence period of dead cysts. Given that in most adult cell types, the M-phase lasts for approximately 1 hour (34–36), this method considerably reduces the required duration of time-lapse imaging. Furthermore, we found that in the *Drosophila* testis, the average number of dead cysts in phase-I was much smaller than that of total cysts (Table S2, S6). Consequently, assuming “zero death” in equation (1) only marginally affects the lifespan estimations (Table S7). This result implies that, if the frequency of death is negligible, then the persistence time will have a much smaller effect on the lifespan estimations, eliminating the requirement of time-lapse measurement of the death progression.

### The rates of germline cells anomalously slow down at the midpoint of transit-amplification

The lifespan estimations indicate a slow down after the 1^st^ TA division and subsequent acceleration that over-compensates the slowdown at the 4-cell stage (Fig. 4B). Termination of *held-out-wing* (How) at the 2-cell stage, coincides with the initial slowdown (17). Moreover, the TA acceleration coincides with that of the increase in bam expression (10) and the termination of the TFGβ signaling gradient (37, 38). How maintains the Cyclin B levels in the *Drosophila* male germline (17), and Bam stabilizes Cyclin A in the *Drosophila* female germline (39). A recent study also suggested that loss of Bam or increased TFGβ signaling slows down the male germline divisions at the 4-cell stage (13). Therefore, we posit that the loss of How expression and the TFGβ signal mediated suppression of bam could prolong the cell cycle length at the 2-cell stage. Recent transcriptomic analysis of single-cysts in the *Drosophila* testis reported a relatively higher level of wee1 (Cdk1 inhibitor) expression in the GBs and high levels of both *cyclinB3* and *twine* (*cdc25*) expression in the 4-cell cysts (40). These latter events coincide with the attenuation of the TGFβ signaling gradient and enhanced bam expression. Presumably, the wee1 expression in GBs would delay the G2/M transition, increasing the cell cycle length of the 2-cell stage; whereas, *cycB3 and twine* expressions in the 4-cell stage would promote the G2/M transition shortening the cell cycle length. What is the significance of these anomalous changes in cellular lifespans? Prolonged interphase at the 2-cell stage probably causes a transition of the cell-fate from a stem-cell mode to a TA mode. In other words, this prolonged interphase at the 2-cell stage could facilitate the transcriptomic changes required for the impending induction of meiosis (Fig. 4C). Alternatively, this pause could be a consequence of the transcriptomic switch.

### Germline cells communicate with their daughters to regulate the rate of TA divisions

Previous studies proposed that the birth of the stem cell progeny sets a molecular clock that counts time before differentiation (10, 41, 42). This theory, however, fails to explain how a stereotypic differentiation clock fine-tunes along with the alterations in rates of TA divisions under different conditions(1–7). Here, we show that altering the lifespans of GSCs and GBs through autonomous perturbations of cell-cycle regulators also modifies the lifespans of the subsequent TA stages (Fig. 4B) and resets the differentiation clock accordingly (Fig. 4C). As a consequence, the extent of germline amplification remains the same; thus, maintaining homeostasis. This conclusion, however, is at variance with the previous conjecture derived by altering the division rates of the late TA stages that induced a premature differentiation at the 8-cell stage or delayed the differentiation till the 32-cell stage (10, 13). One significant difference between the previous experiments and those described above is that we perturbed the rate of the stem cell divisions. The results imply that stem cells can send a forward signal to regulate the division rates of their TA daughters. A similar forward regulation has been reported in *Drosophila* intestinal lining (43) as well as mammalian tracheal epithelia (44). Together, these results suggest that the stem cells communicate with their daughters to regulate the differentiation program and homeostasis.

We envisage that this method can be adapted to estimate lifespan changes in the linages of analogous stem cell systems by marking their mitotic activity with suitable modifications. A limitation of this method is the requirement of stage-wise discrimination of cellular lineage within a tissue. Recent studies have identified markers that separate TA stages from stem cells aiding their identification in various systems (9, 45–48). The method also has a potential application in non-analogous systems in steady-state such as the progression of pathogenic infections or a demographic class in a population.

## Materials and methods

All stocks and crosses were maintained on standard *Drosophila* medium at 25°C. The method used for obtaining the vasa-positive TA stage-wise cyst count (from testes immunostained with anti-vasa and anti-armadillo) has been presented in our previous study (13). The dataset used to compute GSC and TA stage-wise mitotic indices (sum of phosphoHistone-3 positive cysts divided by the sum of vasa-positive cysts at each stage) was presented in this study (13).

### *Ex vivo* imaging of testis

Testes from 4-day old flies were dissected and placed on a poly-lysine coated glass coverslip of a glass-bottom petri dish (P4707; Sigma Chemical Co. USA). For determination of the M-phase duration, the testes were immersed in Schneider’s insect medium (Sigma Chemical Co. USA) and imaged for 2 to 4 hours. The imaging interval was set as 5-6 minutes to minimize phototoxicity; therefore, the estimates have an intrinsic error of ± 5/6 minutes. To estimate the persistence time of dying cysts, dissected testes were incubated in Lysotracker RedDND-99 (ThermoFisher Scientific) in PBS (1:1000 dilution) for 30 minutes and then imaged in PBS for 3 to 4 hours. was re-defined as the phase-I to phase-II transition time, identified by the increase in Lysotracker staining intensity (Fig. S1A) or the shrinkage of cell size (Fig. S1B).

### Image acquisition and analysis

Images were acquired using Olympus FV1000SPD laser scanning confocal microscope using 40X (1.3 NA), or Olympus FV3000SPD laser scanning confocal microscope using 60X (1.42 NA), 40X (1.3 NA), and 10X (0.4 NA) objectives. Multiple optical slices were collected to cover the entire apical part of the testis. Images were analyzed using ImageJ® (http://fiji.sc/Fiji). The Cell-counter^™^ plugin was used for the quantification of the immunostained cysts.

### Statistical analysis

To calculate the variation in the lifespan estimates, the medians of the 1^st^ quartile and 3^rd^ quartile, and the overall median of the data for the M-phase period and vasa-positive counts were used. Student’s T-test was used to calculate P-values unless otherwise mentioned. Origin (OriginLab, Northampton, MA), Graphpad online software (https://www.graphpad.com/), and Microsoft Excel (2013) were used for statistical analyses.

## Author Contributions

PG: Germ cell death distribution in all genotypes and time-lapse experiments to measure the persistence time of dying cysts. KR, DM, PG: time-lapse experiment to measure the M-phase duration. PG, KR: data analysis. NN, PG, SG, and KR: Conceptualization of the mathematical theory. PG and KR: figures composition and manuscript writing.

## Acknowledgments and funding sources

We thank Benny Shilo, Weismann Institute, Israel; Vienna *Drosophila* Resource Center, Austria; and Developmental Studies Hybridoma Bank, Iowa, USA; for fly stocks and other reagents. KR, NN, PG, and SG were supported by the intramural grant of Dept. of Atomic Energy (DAE), Government of India, to TIFR. The study was partly supported by the Department of Biotechnology, Ministry of Science and Technology, India grant (BT/PR/4585/Med/31/155/2012; dated 28 September 2012) and in part by the DAE, TIFR grant 12-R&D-TFR-5.10-100.

## Supplementary Information

### Supplementary Information Text

#### The mathematical modeling of transit-amplifying (TA) divisions in the male germline of *Drosophila melanogaster*

##### A demographic study of germline TA population

This demographic model states that the ratio of numbers of SGs in two stages at any given instant is equal to the time spent in those stages with a correction term due to the germ cell death. This has the useful consequence that for a population in a steady-state, enumerating the numbers of different SGs at any one instant of time (time cross-sectional data) allows us to immediately determine the relationship between the periods that any individual spends in different developmental stages, without recording the entire process of transit amplification.

###### 1. Decomposition of the population into generations

TA divisions in *Drosophila melanogaster* entail the transition of a 1-cell gonialblast to 16-cell spermatogonia (a total of 5 stages) inside an encapsulation formed by two somatic cells; wherein all the individual spermatogonial stages can be easily visualized and quantified. TA comprises a total of four mitotic divisions which are performed by GB, 2-cell, 4-cell, and 8-cell. The 16-cell spermatogonia develop into 16-cell spermatocytes which undergo meiosis. Our analysis is restricted to the four TA divisions.

In our demographic model, we consider that a population *X* is formed by the collection of all SGs in the first four stages of TA from diverse individuals of a particular genetic background. This means that as a set, at any instant of time the population *X*(*t*) decomposes as a disjoint union of subsets corresponding to each phase.

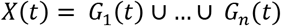

Where period *G_i_*(*t*) is the set of spermatogonia in the stage *i* at the time *t*.

A new individual enters the population as a GB called as a spermatogonial cyst of stage 1, that is, a member of *G*_1_. It successively transits through the generations *G*_2_ (2-cell), *G*_3_ (4-cell) and *G*_4_(8-cell), and it finally exits the population as it enters the 16-cell SG before the meiotic cell cycle.

###### 2. TA stage as a random variable

Let a spermatogonial cyst be picked randomly at time *t* so that any two spermatogonial cysts in the population have an equal chance of getting picked up. This defines the stage of a spermatogonial cyst as a discrete random variable of the population at any time, *i*, which takes four discrete values (GB, 2-cell, 4-cell, 8-cell). We always assume that if randomly drawn from a collection of testes, any two spermatogonial cysts are independent in the sense that, knowledge of the stage of any one of them does not give any additional knowledge of the stage of the other.

###### 3. The probability distribution of the population

Let *p_i_*(*t*) denote the probability for the stage of the SG to be *i* at time where *i* = GB, 2cell, 4cell, 8cell. Note that each *p_i_*(*t*) > 0, and *p*_1_(*t*)+···+ *p*_4_(*t*) = 1. This defines a finite probability distribution which corresponds to the above random variable, *P*(*t*) = (*p_GB_*(*t*),…, *p*_8*cell*_(*t*)). It lies in the three-dimensional convex simplex defined in co-ordinate terms by the equations *x_i_* > 0 for all *i* and *x_GB_* + ··· + *x*_8*cell*_ = 1. As any two SGs have the same chance of being picked up, if denotes the total number of SGs which are in the *i*^th^ stage at time *t*, then, by the basic colored balls in an urn model of probability we must have,

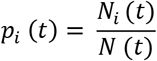

Where *N*(*t*) = *N*_1_(*t*) + ··· *N*_4_(*t*) is the total size of the population, and so we can write

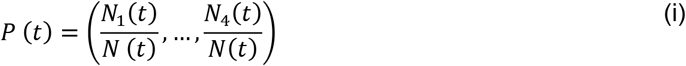

###### 4. The steady-state assumption

The population of interest to us is the collection of all spermatogonial cysts in the first four stages from all the flies belonging to a common genetic background. We will regard this population to be in a steady-state if there are no significant fluctuations in the population over a week. Sampling at 3-time points (namely, 0 days, 4 days, and 8 days post-eclosion of the adult fly) showed that the relative proportions *N_i_*(*t*)/*N*(*t*) of individuals in the first 4 stages remained constant over this period of the life of a juvenile fly (Figure 1D). As the period of the first week in the life of an adult is much smaller than both the lifespan of an individual fly and the period of our experimental observations, this property of the ratios may be described in technical terms as that of being in a quasi-steady state (we will drop the word ‘quasi’ for brevity).

Such a state can arise from the aggregate effect of the behavior of individual testis in flies by a constant rate of input from the germline stem cells and a constant rate of output by the differentiation of 16-cell spermatogonia to 16-cell spermatocytes over the first week of the adult life.

###### 5. Cell cycle lengths/ lifespans of various TA stages

For such a steady-state population belonging to a common genetic background maintained at constant environmental parameters, it can be reasonably assumed that any two spermatogonial cysts spend approximately the same period *T_i_* in the given stage *i*. Of course, the different periods *T_GB_, T*_2*cell*_, *T*_4*cell*_, and *T*_8*cell*_ could have very different values.

Let *T* = *T_GB_* + *T*_2*cell*_ + *T*_4*cell*_ + *T*_8*cell*_ denote the total period (lifespan) that any spermatogonial cyst spends within the population of our interest (namely, the first four stages). To begin with, let us make the simplifying assumption that no deaths occur during the first four stages of a spermatogonial cyst (the deaths are accounted for later). As the mitotic divisions of the germline stem cell are not synchronized, the time of entry of any spermatogonial cyst into the population is uniformly randomly distributed in a cyclic time interval of length *T*. Therefore, under the assumption of no germ cell deaths, as per the uniform continuous probability distribution model, the probability *p_i_* that a randomly chosen SG is in stage *i* at a randomly chosen instant of time, is given by the equation

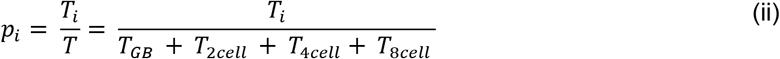

Therefore, equating the two expressions (i) and (ii) for *p_i_* gives us the equalities *p_i_* = *N_i_*/*N* = *T_i_*/*T*, equivalently,

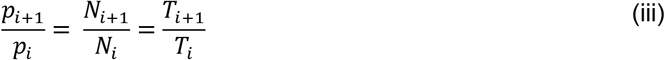

###### 6. Partition of the population into subpopulations

In our model, the population is naturally divided into disjoint subpopulations that reside in distinct testes, so these subpopulations are self-contained from the perspective of their time evolutions within the whole population in the following sense: a spermatogonial cyst in any particular testis spends its entire life-time within that single testis. Each new spermatogonial cyst enters the population by entering a particular testis, goes through all the life stages within that testis, and then exits the testis and the population simultaneously. We will always assume that the stages of individuals within any subpopulation are independent in the sense that for any pair of individuals in a common subpopulation, the knowledge of the stage of the first individual does not give any additional knowledge about the stage of the second individual. It is noteworthy that any two testes contain an approximately equal number of spermatogonial cysts at any chosen common stage (Figure 1D).

###### 7. The necessity for accounting for germ cell deaths

The above set of equations (i, ii, iii) is based on the assumption that the probability of germ cell death during transit amplification is negligible, that is, a 1-cell GB once formed, successfully undergoes the entire transit amplification in 100% of the cases producing a 16-cell SG. However, we observed that a significant proportion of germ cells die during transit amplification (Table S3). This shows that there is a non-zero possibility of germ cell death at each stage of transit amplification. In our model, we have assumed that germ cell death occurs only after the exit of the germ cell from the germline cell cycle at the end of a stage *i* (on successively completing the developmental phases G1, S, G2, and M for stage *i*), and before its possible re-entry into the germline cell cycle in the phase G1 at the beginning of the stage *i* + 1.

###### 8. Survival probability (*s*_*i*+1_)

According to this assumption, after the end of the cell cycle of stage *i*, a spermatogonial cyst either successfully transitions to stage *i* + 1, or undergoes germ cell death. Suppose that on average the transition of *n* individuals from stage *i* to gives rise to *s*_*i*+1_ times *n* individuals in stage *i* + 1. The number *s*_*i*+1_ will be called the survival probability between stages *i* and *i* + 1. If each individual in the stage *i* forms on an average exactly one individual in stage *i* + 1, then we have *s*_*i*+1_ = 1. However, because commonly a certain non-zero proportion of spermatogonial cysts die instead of succeeding in making this transition, we have *s*_*i*+1_ < 1.

Suppose that 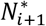 is the hypothetical number of spermatogonial cysts at stage *i* + 1 that would have existed in the population if the survival probability were equal to 1, *s*_*i*+1_, that is, if there were no germ cell deaths during this transition. As seen above we will then get 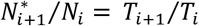. However, the actual number *N*_*i*+1_ is *s*_*i*+1_ times 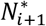. Therefore, a substitution gives (*N*_*i*+1_/*s*_*i*+1_)/*N_i_* = *T*_*i*+1_/*T_i_*, that is,

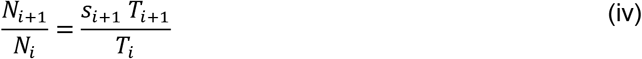

###### 9. Calculation of the survival probability (*s*_*i*+1_) in terms of germ cell death

At any instant of observation, let *D*_*i*+1_ denote the number of dead spermatogonial cysts which have died after the transition from stage *i* to stage *i* + 1, before entering the germline cell cycle of stage *i* + 1. Let *A*_*i*+1_ denote the period for which any dead spermatogonial cyst, that has died at the end of the transition from stage *i* to *i* + 1, remains visible in the testis. We may regard all such dead spermatogonial cysts as forming a new stage (*i* + 1)^*^, which persists for a duration of *A*_*i*+1_. The probability of transition from stage *i* to stage (*i* + 1)^*^ is *d*_*i*+1_ = 1 – *s*_*i*+1_. Therefore, the equality (iv) above, with *D*_*i*+1_ in place of *N*_*i*+1_, *A*_*i*+1_ in place of *T*_*i*+1_ and *d*_*i*+1_ in place of *s*_*i*+1_ now gives *D*_*i*+1_/*N_i_* = *d*_*i*+1_ * *A*_*i*+1_/*T_i_*, that is,

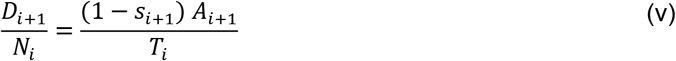

Rearrangement gives

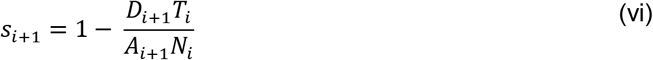

###### 10. Solving the equations to obtain the lifespans

The values of *N_i_* were obtained by enumerating the number of spermatogonial cysts (Table S2). Inserting these values, (iv) and (vi) give two equations in the three unknowns *T_i_, T*_*i*+1_ and *s*_*i*+1_.

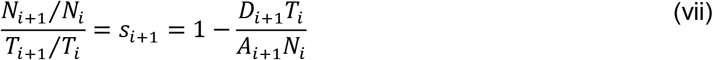

Eliminating *s*_*i*+1_ and solving for *T*_*i*+1_ gives

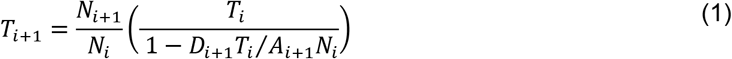

Using the above equation iteratively for *i* = 1, 2, 3, the periods *T*_2_, *T*_3_ and *T*_4_ are determined in terms of the period *T*_1_. We call this method based on equation (1) the “sequential estimation” method. It requires the data (*N_i_, N*_*i*+1_, *D*_*i*+1_, *A*_*i*+1_, *T_i_*) of two successive spermatogonial stages namely, stage *i* and *i* + 1.

Alternatively, one can divide the spermatogonial cyst population of stage *i* into two sub-stages: the cyst population before the M-phase and in M-phase. We denote these two stages as stage *i′* and *i″*, respectively. As mentioned earlier, our assumption implies that no deaths can occur during the phases G1, S, G2, and M of the germline cell cycle itself. As the sub-stage *i′* is by definition made up of the phases G1, S, and G2, and as the substage is just the corresponding phase M, the assumption implies that any spermatogonial cyst in sub-stage *′* has a 100% probability of transitioning to the sub-stage *i″*. Consequently, we have,

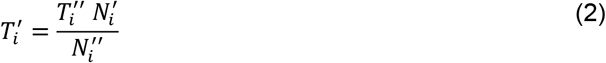

We call this method based on equation (2) the “individual estimation” method. Here 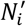, and 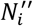 are the number of vasa-positive, phosphorylated-histone3-negative spermatogonial cysts and vasa-positive, phosphorylated-histone3-positive spermatogonial cysts in stage *i*, respectively. 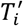, and 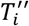 are the durations of stages *i′* and *i″*, respectively.

###### 11. Calculation of *T*_4_ from the lifespan of GSCs (*T*_0_)

Consistent with earlier studies (49), we did not find any dead GSCs in any of the genetic backgrounds examined in this study. Therefore, substituting *D_i_* = 0 in the equation (1) with *i* = GSC gives the duration spent by the GSC in stage *i′* by the formula,

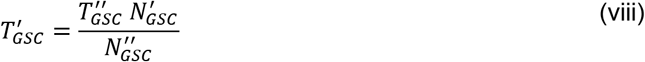

We already know of the period 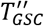 as shown in Figure 3. 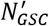, and 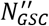 can be determined by the enumeration of vasa-positive, phosphorylated-histone3-negative GSCs and vasa-positive, phosphorylated-histone3-positive GSCs, respectively. As mentioned earlier, the lifespan of the GSC can be obtained as 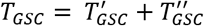.

A previous study reported that the GSCs undergo symmetric differentiation (13%) and that of the symmetric renewals (7%), with low frequency (18). If these proportions are correct, then on average per every 100 divisions of GSCs, 2 × 7 + 80 = 94 GSCs will form while 6 GSCs will be lost. The inter-division time for a GSC is reported as 14 hours (50). Thus, the total number of divisions performed by a single GSC in 8 days (192 hours) will be equal to *n* = 192 / 14 = 13.71 ≈ 14. Therefore, if we begin with a certain number of GSCs on the day of the fly eclosion (0-day), after 8-days we will have (94/100)^14^ = 42% of the starting number of GSCs. However, in contrast to this conclusion our observation indicated that upon quantification of GSCs at 0-day, 4-day, and 8-days post adult eclosion, we found that the GSC number does not vary significantly (Figure 1E).

Therefore, one may infer that, at least for the first 8-days of the adult fly life, the percentage of symmetric differentiation should be equal to the percentage of symmetric renewals among all GSC divisions. Let us denote the common probability of symmetric differentiation and symmetric renewal as *p* and that of the asymmetric division as *q* so that we have 2*p* + *q* = 1. This implies that only q proportion of GSCs form GBs. The GBs so formed either successfully re-enter the cell cycle at stage 1 (with probability *s_GB_*) or die (with probability 1 – *s_GB_*). Therefore, the final proportion of GSCs forming GBs will be given by the product 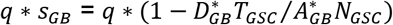. Hence arguing as before, we have

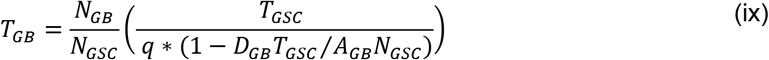

A GSC can form a 2-cell spermatogonial cyst directly by symmetric differentiation or by an asymmetric division forming a GB, followed by a GB division. The probability of the direct formation of a 2-cell stage from a GB is the product of the probability of the asymmetric division (equal to *p*) and the survival probability of the 2-cell stage (equal to *s*_2*cell*_). Since the probability of a GSC forming a GB is 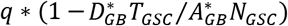, the probability of a GB forming 2-cell spermatogonial stage will be, *q* * *S_GB_* * *s*_2*cell*_, where *s*_2*cell*_ is the survival probability of a 2-cell stage. Thus, the probability of a GSC forming a 2-cell spermatogonial cyst (either directly or by passage through a GB) is *q* * *s_GB_ * s*_2*cell*_ + *p* * *s*_2*cell*_, or simply (*q * s_GB_ + p*) * *s*_2*cell*_. This gives us

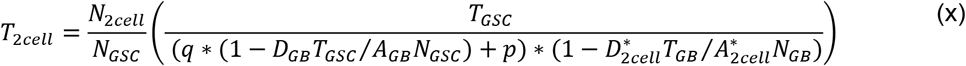

To determine the periods *T*_3_ and *T*_4_ in terms of the period *T*_2_ using the sequential method, equation (2) can be used iteratively for *i* = 4cell, 8cell. For our study, we have assumed that the frequencies of symmetric differentiation and symmetric renewals are negligible as compared to the frequency of asymmetric division. However, we have calculated the lifespans in the control background including the reported values for symmetric differentiation and symmetric renewals in wild-type (Figure S4).

###### 12. Determination of the clearance time of a dying SG (A) and GCD

The germline death progression could be divided into four successive phases based on the staining pattern of a nuclear stain (DAPI), Lysotracker, Lamin, and the germline marker - Vasa (22). The first two phases are identifiable and can be classified into GB, 2-cell, 4-cell, etc. depending on the anti-vasa and Lysotracker staining. Therefore, we picked the time spent by the dying cell in phase-I as the time of clearance of a dead spermatogonial cyst. Note that one may apply the formula in equations (1) with *D*_*i*+1_ as the average number of spermatogonial cysts of stage*i* + 1 in phase-I of death and *A*_*i*+1_ as the time spent in phase-I of death.

**Fig. S1.**
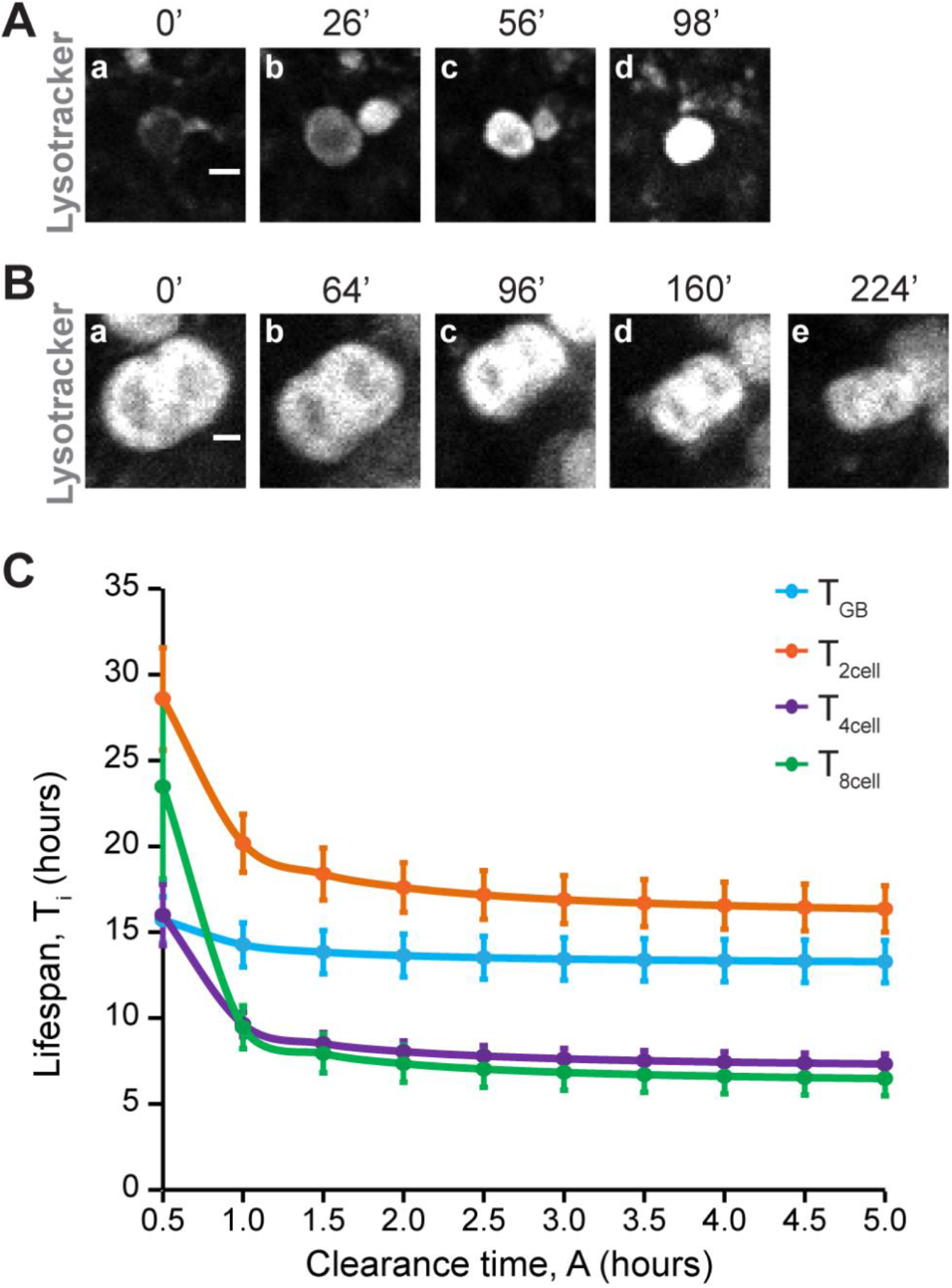
Clearance time of dead cyst A) Montage of a GB transition from phase-I (a, b) to phase-II (c) of germ cell death, demonstrating an increase in the Lysotracker intensity. B) Montage of a 2-cell cyst transition from phase-I (a, b, c) to phase-II (e) of germ cell death, demonstrating size reduction. Time intervals in minutes are indicated at top of the panels. (Scale bar ~5μm). C) Line plot shows predicted lifespans of GB (***T_GB_***, Blue), 2-cell (***T*_2*cell*_**, Orange), 4-cell (***T*_4*cell*_**, Purple), and 8-cell (***T*_8*cell*_**, Green) in control (*nosGal4vp16>UAS-EGFP*) background with the persistence time (***A***) ranging from 0.5 hours to 5 hours.

**Fig. S2.**
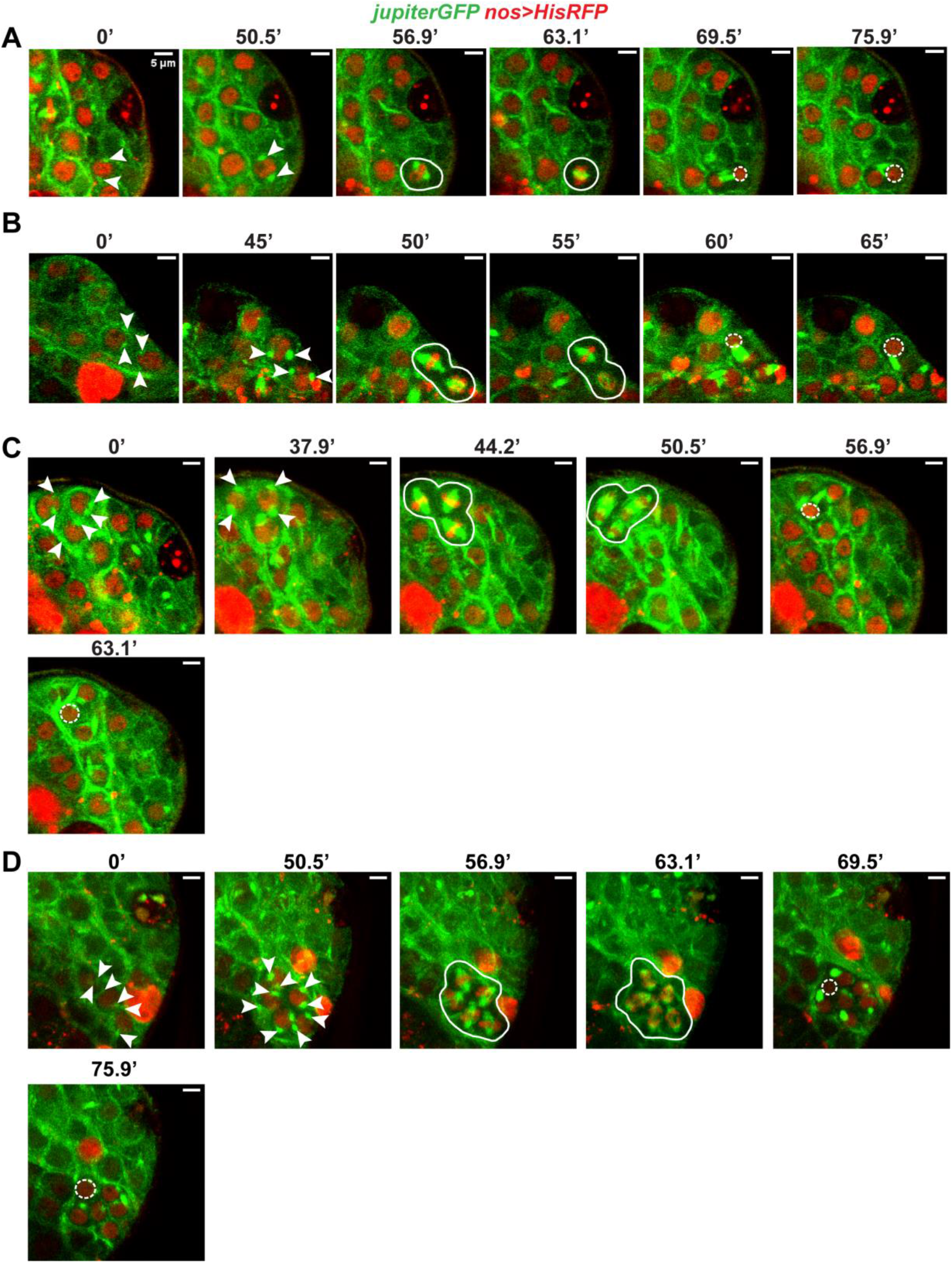
Montage of a time-lapse image of a GB (A), 2-cell (B), 4-cell (C) and 8-cell (D) (*JupiterGFP* (Green), *nos>HisRFP* (Red)), undergoing M-phase. Arrowheads mark the position of the separated centrosomes. White circles label the cells visible in the plane of imaging. White dashed circles depict the increase in the nuclear size marking the end of telophase. Time intervals in minutes have been indicated on top of the panels. (Scale bars ~ 5μm).

**Fig. S3.**
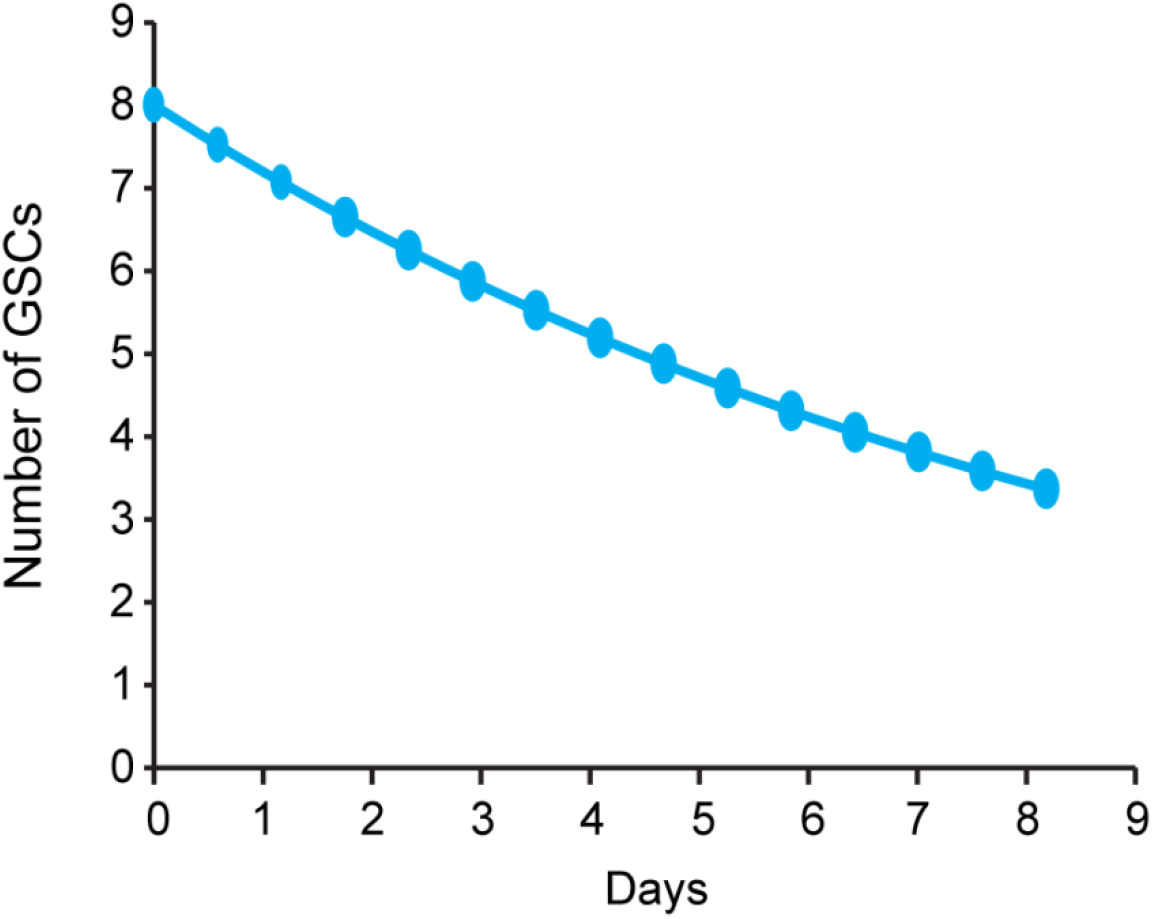
Line plot shows the predicted decline in GSC numbers with days after adult fly eclosion calculated by considering % Asymmetric division = 80%, % Symmetric renewals = 7% and % Symmetric differentiation = 13% (Reported in (18)).

**Fig. S4.**
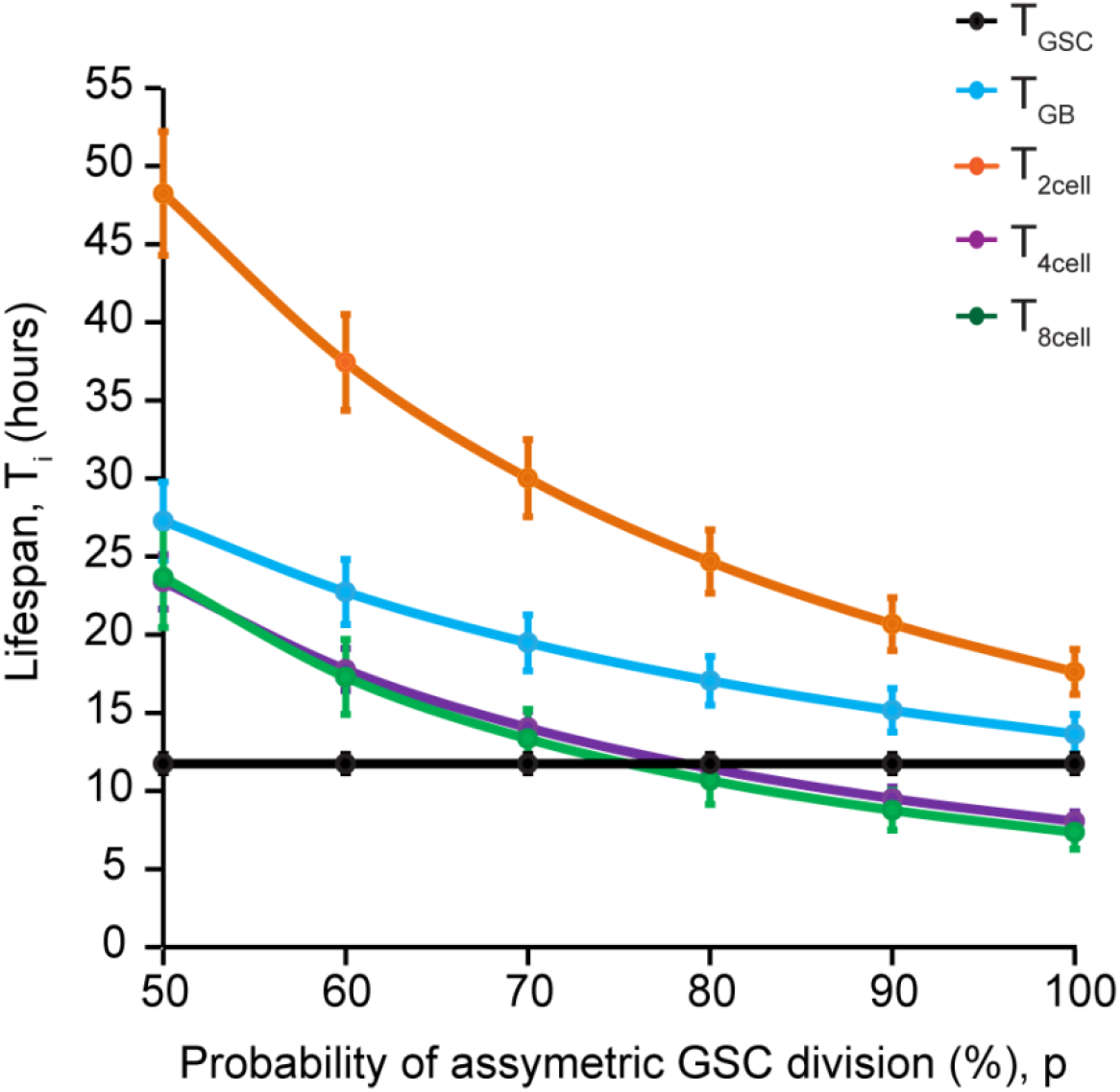
The line plot shows the predicted lifespans of GSC (***T_GSC_***, Black), GB (***T_GB_***, Blue), 2-cell (***T*_2*cell*_**, Orange), 4-cell (***T*_4*cell*_**, Purple), and 8-cell (***T*_8*cell*_**, Green) in control (*nosGal4vp16>UAS-EGFP*) background with the % probability of asymmetric division ranging from 1 to 0.5 (q = 100, 90, 80, 70, 60, 50).

**Table S1:** Stage-wise Mitotic index in control and cell cycle perturbed backgrounds.

| Genotype | Mitotic index |  |  |  |  |
| --- | --- | --- | --- | --- | --- |
|  | GSC | GB | 2-cell | 4-cell | 8-cell |
| ***nos*> control (40)** | 0.0962 | 0.0933 | 0.0639 | 0.1264 | 0.1373 |
| ***nos*>*cycE*<sup>dsRNA</sup> (24)** | 0.0482* | 0.0629 | 0.0448 | 0.0870 | 0.0505 |
| ***nos*>*Cdk1*<sup>dsRNA</sup> (21)** | 0.0417† | 0.0455 | 0.0500 | 0.0714 | 0.0909 |
| ***nos*>*stg*<sup>dsRNA</sup> (23)** | 0.0667 | 0.1074 | 0.0493 | 0.0787 | 0.1176 |
| ***nos*>*stg* (18)** | 0.1835‡ | 0.2024§ | 0.1045 | 0.1707 | 0.1333 |

\* Fisher's exact test, $P = 0.0476$
† Fisher's exact test, $P = 0.0312$
‡ Fisher's exact test, $P = 0.0116$
§ Fisher's exact test, $P = 0.0038$

**Table S2:** Stage-wise cyst distribution profile (Median ± Interquartile range)

| **Genotype** | **GSC** | **GB** | **2-cell** | **4-cell** | **8-cell** |
| --- | --- | --- | --- | --- | --- |
| ***nos*> control (40)** | 8 ± 1.25 | 9 ± 3 | 10 ± 3 | 5 ± 1.25 | 4 ± 2 |
| ***nos*>*cycE*<sup>dsRNA</sup> (25)** | 9<sup>†</sup> ± 2 | 7<sup>†</sup> ± 2 | 9* ± 2 | 4 ± 2 | 4 ± 2 |
| ***nos*>*Cdk1*<sup>dsRNA</sup> (21)** | 7 ± 2 | 6<sup>‡</sup> ± 2 | 8<sup>†</sup> ± 2 | 4 ± 2 | 4 ± 2 |
| ***nos*>*stg*<sup>dsRNA</sup> (23)** | 7<sup>†</sup> ± 1 | 5<sup>‡</sup> ± 2 | 6<sup>‡</sup> ± 3 | 4 ± 2 | 3* ± 1.5 |
| ***nos*>*stg* (18)** | 9* ± 2 | 9.5 ± 1 | 12<sup>†</sup> ± 2.75 | 7<sup>‡</sup> ± 2 | 7<sup>‡</sup> ± 3 |

\* Mann-Whitney-U test, $P < 0.05$
† Mann-Whitney-U test, $P < 0.01$
‡ Mann-Whitney-U test, $P < 0.001$

**Table S3:** Average number of cysts in Phase-I of Germ cell death (*D*_*i*+1_) in different genetic backgrounds.

| Genotype | Average cell death (PHASE-I) |  |  |  |  |
| --- | --- | --- | --- | --- | --- |
|  | GSC | GB | 2-cell | 4-cell | 8-cell |
| *nos*> control (54) | 0 | 0.06 | 0.09 | 0.04 | 0.06 |
| *nos*>*cycE*<sup>dsRNA</sup> (13) | 0 | 0.08 | 0.15 | 0 | 0.15 |
| *nos*>*Cdk1*<sup>dsRNA</sup> (29) | 0 | 0 | 0 | 0 | 0.03 |
| *nos*>*stg*<sup>dsRNA</sup> (21) | 0 | 0 | 0 | 0.05 | 0.1 |
| *nos*>*stg* (10) | 0 | 0 | 0.4 | 0 | 0 |

**Table S4.** Clearance time (A) recorded for different TA stages in various control genetic backgrounds.

| Genotype | TA stage | Phase-I persistence time (Hours) |
| --- | --- | --- |
| *tj>* | Gonialblast | >1.6 |
|  | Gonialblast | 1.07* |
|  | Gonialblast | >0.55 |
|  | Gonialblast | >0.95 |
|  | 2-cell | >3.73 |
|  | 2-cell | 0.9* |
|  | 8-cell | >3.73 |
|  | 8-cell | >1.27 |
|  | 16-cell | >3.73 |
|  | 16-cell | >2.20 |

The total number of time-lapse images acquired from the genetic controls were 71 - *tjGal4* (total 57) and *nosGal4* (total 14).
\* Both the onset of Phase-I and transition to Phase-II were recorded

**Table S5:**
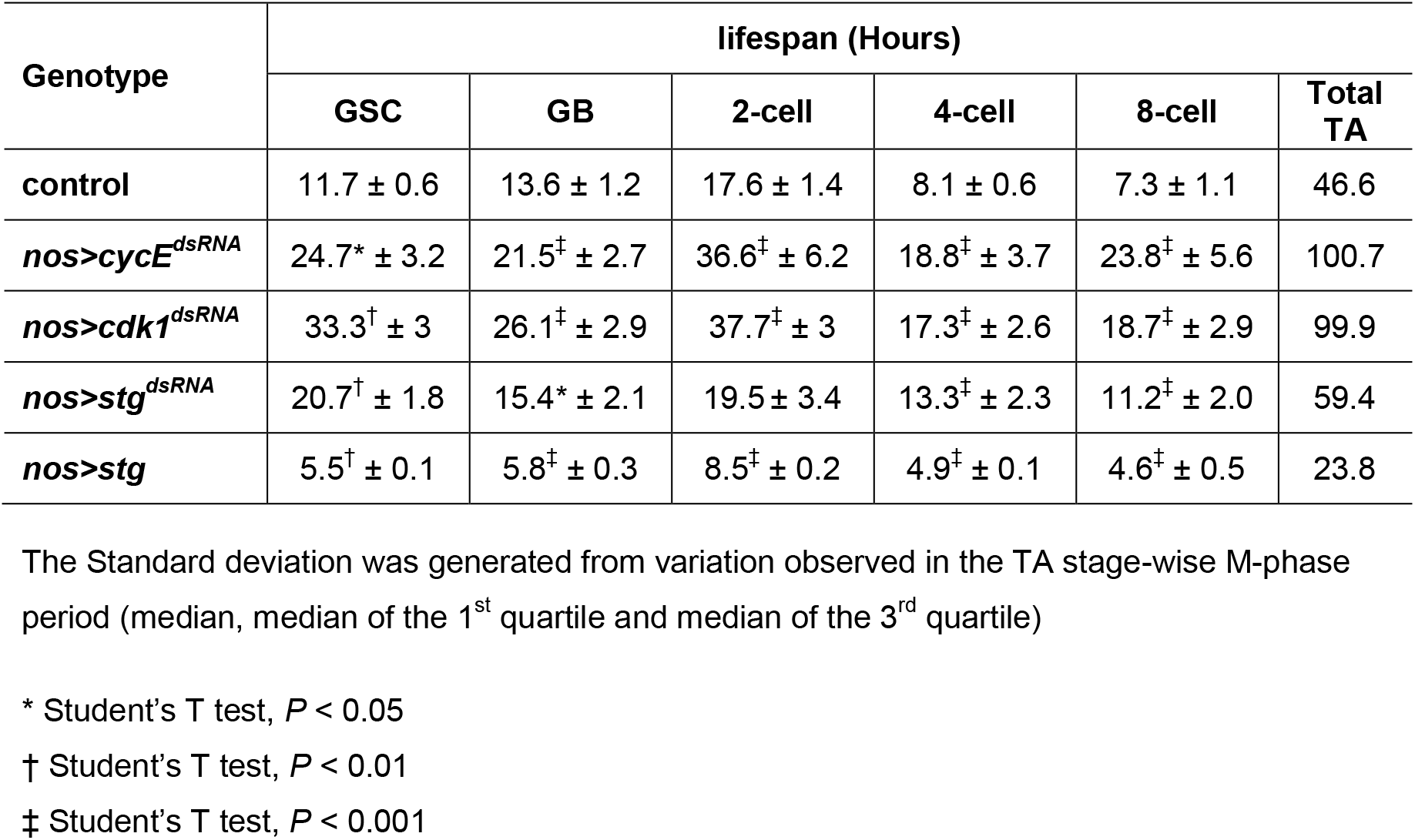
Lifespan estimations (Average ± SD) using the equation (1) in different genetic backgrounds (Presented in Fig. 3)

**Table S6:**
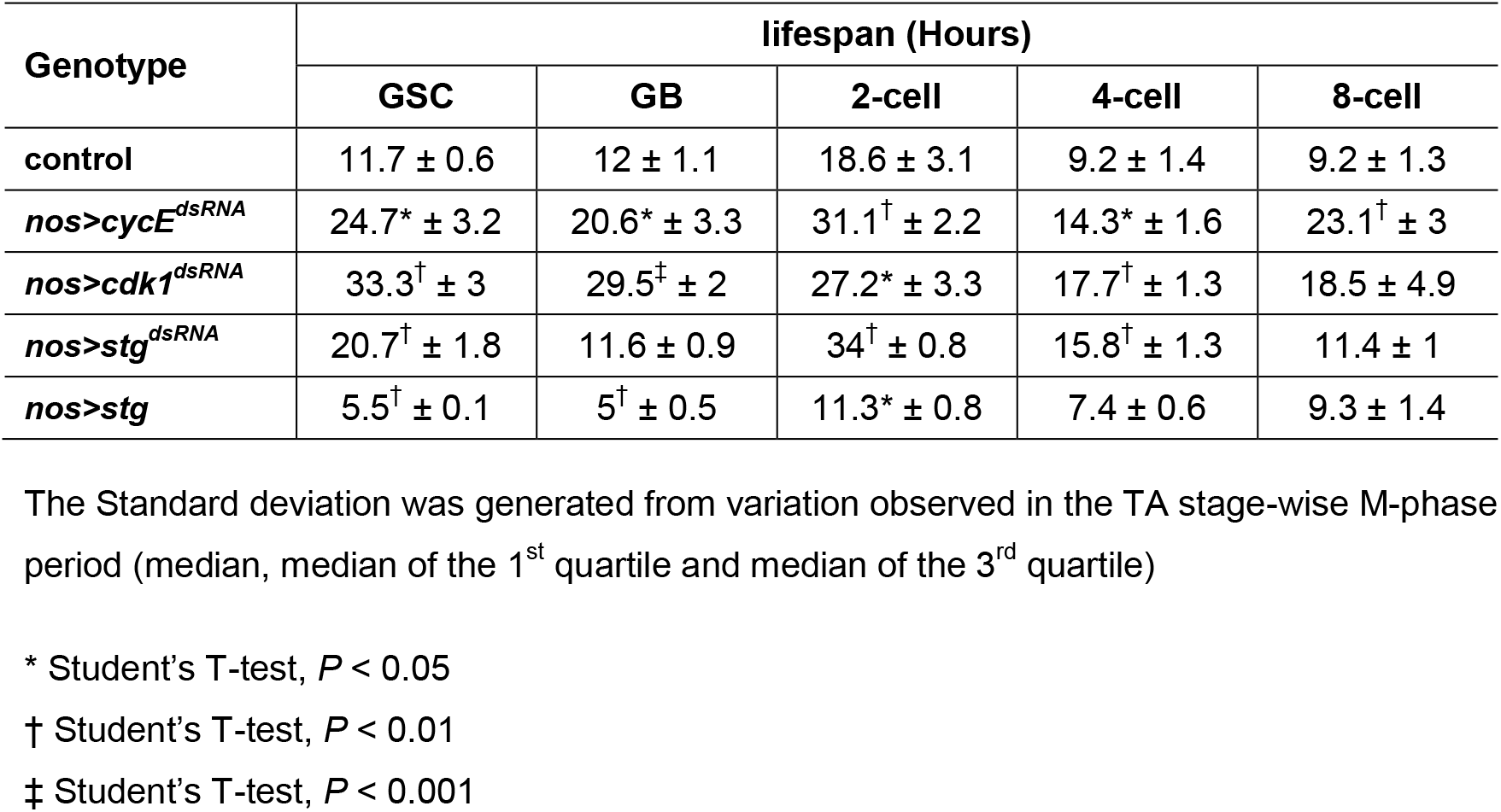
Lifespan estimations (Average ± SD) using the equation (2) in different genetic backgrounds.

| Genotype | lifespan (Hours) |  |  |  |  |
| --- | --- | --- | --- | --- | --- |
|  | GSC | GB | 2-cell | 4-cell | 8-cell |
| control | 11.7 ± 0.6 | 12 ± 1.1 | 18.6 ± 3.1 | 9.2 ± 1.4 | 9.2 ± 1.3 |
| *nos>cycE<sup>dsRNA</sup>* | 24.7* ± 3.2 | 20.6* ± 3.3 | 31.1<sup>†</sup> ± 2.2 | 14.3* ± 1.6 | 23.1<sup>†</sup> ± 3 |
| *nos>cdk1<sup>dsRNA</sup>* | 33.3<sup>†</sup> ± 3 | 29.5<sup>‡</sup> ± 2 | 27.2* ± 3.3 | 17.7<sup>†</sup> ± 1.3 | 18.5 ± 4.9 |
| *nos>stg<sup>dsRNA</sup>* | 20.7<sup>†</sup> ± 1.8 | 11.6 ± 0.9 | 34<sup>†</sup> ± 0.8 | 15.8<sup>†</sup> ± 1.3 | 11.4 ± 1 |
| *nos>stg* | 5.5<sup>†</sup> ± 0.1 | 5<sup>†</sup> ± 0.5 | 11.3* ± 0.8 | 7.4 ± 0.6 | 9.3 ± 1.4 |

The Standard deviation was generated from variation observed in the TA stage-wise M-phase period (median, median of the 1<sup>st</sup> quartile and median of the 3<sup>rd</sup> quartile)
\* Student's T-test, $P < 0.05$
† Student's T-test, $P < 0.01$
‡ Student's T-test, $P < 0.001$

**Table S7:** Lifespans estimates considering no germ cell death (GCD)

| Method | lifespan (Hours) |  |  |  |  |
| --- | --- | --- | --- | --- | --- |
|  | GSC | GB | 2-cell | 4-cell | 8-cell |
| **Sequential ($D_{i+1} = 0$)** | 11.7 ± 0.6 | 13.1 ± 1.2 | 15.6 ± 1.3 | 6.9 ± 0.6 | 6 ± 0.9 |
| **Sequential ($D_{i+1} = \text{observed}$)** | 11.7 ± 0.6 | 13.6 ± 1.2 | 17.6<sup>†</sup> ± 1.4 | 8.1<sup>‡</sup> ± 0.6 | 7.3* ± 1.1 |

The Standard deviation was generated from variation observed in the TA stage-wise M-phase period and vasa-positive cyst counts (median, median of the 1<sup>st</sup> quartile and median of the 3<sup>rd</sup> quartile)
\* Student's T-test, $P < 0.05$
† Student's T-test, $P < 0.01$
‡ Student's T-test, $P < 0.001$

## Notes

### Competing Interest Statement

The authors have declared no competing interest.

## References

1. R. R. Giraddi, et al., Stem and progenitor cell division kinetics during postnatal mouse mammary gland development. Nat. Commun. 6, 1–12 (2015).

2. G. B. Di Gregorio, et al., Attenuation of the self-renewal of transit-amplifying osteoblast progenitors in the murine bone marrow by 17β-estradiol. J. Clin. Invest. 107, 803–812 (2001).

3. M. S. Lehrer, T. T. Sun, R. M. Lavker, Strategies of epithelial repair: modulation of stem cell and transit amplifying cell proliferation. J. Cell Sci. 111 (Pt 1, 2867–75 (1998).

4. R. Ichijo, et al., Tbx3-dependent amplifying stem cell progeny drives interfollicular epidermal expansion during pregnancy and regeneration. Nat. Commun. 8, 1–12 (2017).

5. A. Charruyer, et al., Transit-amplifying cell frequency and cell cycle kinetics are altered in aged epidermis. J. Invest. Dermatol. 129, 2574–2583 (2009).

6. C. Tomasetti, et al., Cell division rates decrease with age, providing a potential explanation for the age-dependent deceleration in cancer incidence. Proc. Natl. Acad. Sci. U. S. A. 116, 20482–20488 (2019).

7. G. N. Paliouras, et al., Mammalian target of rapamycin signaling is a key regulator of the transit-amplifying progenitor pool in the adult and aging forebrain. J. Neurosci. 32, 15012–15026 (2012).

8. H. Li, et al., Rate of Progression through a Continuum of Transit-Amplifying Progenitor Cell States Regulates Blood Cell Production. Dev. Cell 49, 118–129.e7 (2019).

9. O. A. Bayraktar, C. Q. Doe, Combinatorial temporal patterning in progenitors expands neural diversity. Nature 498, 449–455 (2013).

10. M. L. Insco, A. Leon, C. H. Tam, D. M. McKearin, M. T. Fuller, Accumulation of a differentiation regulator specifies transit amplifying division number in an adult stem cell lineage. Proc. Natl. Acad. Sci. U. S. A. 106, 22311–22316 (2009).

11. B. Zhang, Y. C. Hsu, Emerging roles of transit-amplifying cells in tissue regeneration and cancer. Wiley Interdiscip. Rev. Dev. Biol. 6, 1–14 (2017).

12. M. F. Clarke, M. Fuller, Stem Cells and Cancer: Two Faces of Eve. Cell 124, 1111–1115 (2006).

13. P. Gadre, S. Chatterjee, B. Varshney, K. Ray, Cyclin E and Cdk1 regulate the termination of germline transit-amplification process in Drosophila testis. Cell Cycle 19, 1786–1803 (2020).

14. C. Y. Li, Z. Guo, Z. Wang, TGF?? receptor saxophone non-autonomously regulates germline proliferation in a Smox/dSmad2-dependent manner in Drosophila testis. Dev. Biol. 309, 70–77 (2007).

15. B. B. Parrott, A. Hudson, R. Brady, C. Schulz, Control of Germline stem cell division frequency - A novel, developmentally regulated role for Epidermal Growth Factor signaling. PLoS One 7 (2012).

16. S. Gupta, B. Varshney, S. Chatterjee, K. Ray, Somatic ERK activation during transit amplification is essential for maintaining the synchrony of germline divisions in Drosophila testis (2018) https:/doi.org/10.6084/m9.

17. A. C. Monk, et al., HOW Is Required for Stem Cell Maintenance in the Drosophila Testis and for the Onset of Transit-Amplifying Divisions. Cell Stem Cell 6, 348–360 (2010).

18. X. R. Sheng, E. Matunis, Live imaging of the Drosophila spermatogonial stem cell niche reveals novel mechanisms regulating germline stem cell output. Development 138, 3367–76 (2011).

19. K. F. Lenhart, S. DiNardo, Somatic Cell Encystment Promotes Abscission in Germline Stem Cells following a Regulated Block in Cytokinesis. Dev. Cell 34, 192–205 (2015).

20. K. Yacobi-Sharon, Y. Namdar, E. Arama, Alternative germ cell death pathway in drosophila involves HtrA2/Omi, lysosomes, and a caspase-9 counterpart. Dev. Cell 25, 29–42 (2013).

21. H. Yang, Y. M. Yamashita, The regulated elimination of transit-amplifying cells preserves tissue homeostasis during protein starvation in Drosophila testis. Development 142, 1756–1766 (2015).

22. A. C.-Y. Chiang, H. Yang, Y. M. Yamashita, spict, a cyst cell-specific gene, regulates starvation- induced spermatogonial cell death in the Drosophila testis. Sci. Rep. 7, 40245 (2017).

23. R. Giet, D. M. Glover, Drosophila aurora B kinase is required for histone H3 phosphorylation and condensin recruitment during chromosome condensation and to organize the central spindle during cytokinesis. J. Cell Biol. 152, 669–681 (2001).

24. N. Karpova, Y. Bobinnec, S. Fouix, P. Huitorel, A. Debec, Jupiter, a newDrosophila protein associated with microtubules. Cell Motil. Cytoskeleton 63, 301–312 (2006).

25. K. H. Siller, M. Serr, R. Steward, T. S. Hays, C. Q. Doe, Live imaging of Drosophila brain neuroblasts reveals a role for Lis1/dynactin in spindle assembly and mitotic checkpoint control. Mol. Biol. Cell 16, 5127–40 (2005).

26. M. S. Savoian, C. L. Rieder, Mitosis in primary cultures of Drosophila melanogaster larval neuroblasts. J. Cell Sci. 115, 3061–72 (2002).

27. S. Hasan, P. Hétié, E. L. Matunis, Niche signaling promotes stem cell survival in the Drosophila testis via the JAK-STAT target DIAP1. Dev. Biol. 404 (2015).

28. Lindsley, L., D., Tokuyasu K., T., “Spermatogenesis” in The Genetics and Biology of Drosophila. Ashburner, M. and Wright, T.R.F., Eds., (Academic Press 1980) 2, pp. 225–294.

29. T. D. Hinnant, A. A. Alvarez, E. T. Ables, Temporal remodeling of the cell cycle accompanies differentiation in the Drosophila germline. Dev. Biol. 429, 118–131 (2017).

30. C. C. F. Homem, I. Reichardt, C. Berger, T. Lendl, J. A. Knoblich, Long-Term Live Cell Imaging and Automated 4D Analysis of Drosophila Neuroblast Lineages. PLoS One 8, 79588 (2013).

31. J. A. Knoblich, et al., Cyclin E controls S phase progression and its down-regulation during Drosophila embryogenesis is required for the arrest of cell proliferation. Cell 77, 107–20 (1994).

32. B. A. Edgar, P. H. O’Farrell, Genetic control of cell division patterns in the Drosophila embryo. Cell 57, 177–87 (1989).

33. M. Inaba, H. Yuan, Y. M. Yamashita, String (Cdc25) regulates stem cell maintenance, proliferation and aging in Drosophila testis. Development 138, 5079–5086 (2011).

34. A. B. Georgi, P. T. Stukenberg, M. W. Kirschner, Timing of events in mitosis. Curr. Biol. 12, 105–114 (2002).

35. J. Padgett, S. D. M. Santos, From clocks to dominoes: lessons on cell cycle remodelling from embryonic stem cells. FEBS Lett. 594, 2031–2045 (2020).

36. F. D. Sigoillot, et al., A Time-Series Method for Automated Measurement of Changes in Mitotic and Interphase Duration from Time-Lapse Movies. PLoS One 6, e25511 (2011).

37. L. Xia, et al., The fused/smurf complex controls the fate of drosophila germline stem cells by generating a gradient bmp response. Cell 143, 978–990 (2010).

38. E. Kawase, Gbb/Bmp signaling is essential for maintaining germline stem cells and for repressing bam transcription in the Drosophila testis. Development 131, 1365–1375 (2004).

39. S. Ji, et al., Bam-dependent deubiquitinase complex can disrupt germ-line stem cell maintenance by targeting cyclin A. Proc. Natl. Acad. Sci. 114, 6316–6321 (2017).

40. Z. Shi, et al., Single-cyst transcriptome analysis of Drosophila male germline stem cell lineage. Dev. 147 (2020).

41. O. Cinquin, S. L. Crittenden, D. E. Morgan, J. Kimble, Progression from a stem cell-like state to early differentiation in the C. elegans germ line. Proc. Natl. Acad. Sci. U. S. A. 107, 2048–2053 (2010).

42. F.-B. Gao, B. Durand, M. Raff, Oligodendrocyte precursor cells count time but not cell divisions before differentiation. Curr. Biol. 7, 152–155 (1997).

43. B. Ohlstein, A. Spradling, Multipotent Drosophila intestinal stem cells specify daughter cell fates by differential notch signaling. Science (80-.). 315, 988–992 (2007).

44. A. Pardo-Saganta, et al., Parent stem cells can serve as niches for their daughter cells. Nature 523, 597–601 (2015).

45. Y. Li, N. T. Minor, J. K. Park, D. M. McKearin, J. Z. Maines, Bam and Bgcn antagonize Nanos- dependent germ-line stem cell maintenance. Proc. Natl. Acad. Sci. U. S. A. 106, 9304–9 (2009).

46. H. Li, et al., Rate of Progression through a Continuum of Transit-Amplifying Progenitor Cell States Regulates Blood Cell Production. Dev. Cell 49, 118–129.e7 (2019).

47. T. H. Kim, et al., Single-Cell Transcript Profiles Reveal Multilineage Priming in Early Progenitors Derived from Lgr5+ Intestinal Stem Cells. Cell Rep. 16, 2053–2060 (2016).

48. F. Doetsch, L. Petreanu, I. Caille, J. M. Garcia-Verdugo, A. Alvarez-Buylla, EGF converts transit-amplifying neurogenic precursors in the adult brain into multipotent stem cells. Neuron 36, 1021–1034 (2002).

## SI References

1. K. Yacobi-Sharon, Y. Namdar, E. Arama, Alternative Germ Cell Death Pathway in *Drosophila* Involves HtrA2/Omi, Lysosomes, and a Caspase-9 Counterpart. Dev. Cell 25, 29–42 (2013).

2. X. R. Sheng, E. Matunis, Live imaging of the *Drosophila* spermatogonial stem cell niche reveals novel mechanisms regulating germline stem cell output. Development 138, 3367–76 (2011).

3. K. F. Lenhart, S. DiNardo, Somatic Cell Encystment Promotes Abscission in Germline Stem Cells following a Regulated Block in Cytokinesis. Dev. Cell 34, 192–205 (2015).

4. A. C.-Y. Chiang, H. Yang, Y. M. Yamashita, spict, a cyst cell-specific gene, regulates starvation-induced spermatogonial cell death in the *Drosophila* testis. Sci. Rep. 7, 40245 (2017).

